# Learning function from structure in neuromorphic networks

**DOI:** 10.1101/2020.11.10.350876

**Authors:** Laura E. Suárez, Blake A. Richards, Guillaume Lajoie, Bratislav Misic

## Abstract

The connection patterns of neural circuits in the brain form a complex network. Collective signaling within the network manifests as patterned neural activity, and is thought to support human cognition and adaptive behavior. Recent technological advances permit macro-scale reconstructions of biological brain networks. These maps, termed connectomes, display multiple non-random architectural features, including heavy-tailed degree distributions, segregated communities and a densely interconnected core. Yet, how computation and functional specialization emerge from network architecture remains unknown. Here we reconstruct human brain connectomes using *in vivo* diffusion-weighted imaging, and use reservoir computing to implement these connectomes as artificial neural networks. We then train these neuromorphic networks to learn a cognitive task. We show that biologically realistic neural architectures perform optimally when they display critical dynamics. We find that performance is driven by network topology, and that the modular organization of large-scale functional systems is computationally relevant. Throughout, we observe a prominent interaction between network structure and dynamics, such that the same underlying architecture can support a wide range of learning capacities across dynamical regimes. This work opens new opportunities to discover how the network organization of the brain optimizes cognitive capacity, conceptually bridging neuroscience and artificial intelligence.

## INTRODUCTION

The brain is a complex network of anatomically connected and functionally interacting neuronal populations. The wiring of the network allows its components to collectively transform signals representing internal states and external stimuli. Recent technological and analytic advances provide the opportunity to comprehensively map, image and trace connection patterns of nervous systems in multiple species [61, 146], yielding high-resolution “connectomes” of individual brains [10, 130]. Numerous reports have found evidence of non-random topological attributes that theoretically enhance the capacity of the network to process information [146], including high clustering and short path length [8, 65, 131, 158], specialized segregated communities [15, 20, 27, 52, 79, 106], heavy-tailed degree distributions [48, 129], and a core of densely inter-connected hub nodes [143, 147, 164]. The wiring patterns needed to support these attributes entail energetic and metabolic cost, yielding an economic trade-off between efficient information transmission and minimal wiring cost [23]. Despite these insights, the link between macroscale connectivity structure and the computational properties that emerge from network activity is not well understood.

How does the organization of the brain confer computational capacity? One way to address this question is to relate structural connectivity to neural dynamics and emergent patterns of functional connectivity [22, 119, 135, 144]. Another is to relate structural connectivity to individual differences in behaviour [90, 96, 117]. Although both paradigms inform us about functional consequences of network organization, they do not explicitly consider how network organization supports information processing [64]. An alternative way to conceptualize structure-function relationships is to directly consider function as a computational property. Modern artificial intelligence algorithms offer new ways to link structure and function, by conceptualizing function as a computational property [108].

A prominent paradigm to understand how artificial recurrent neural networks extract information from a continuous stream of external stimuli is reservoir computing [80]. Also known as *echo-state networks* or *liquid state machines* [62, 82], this computational schema typically uses randomly connected recurrent neural networks, but arbitrary network architectures can also be used. The networks are trained in a supervised manner to learn representations of external time-varying stimuli, and can be adapted to a wide range of tasks, including speech recognition [124, 151, 152], motor learning [60, 78, 113], natural language processing [53, 142], working memory [101, 133] and spatial navigation [3, 136]. A significant benefit of the reservoir framework is that arbitrary dynamics can be super-imposed on the network, providing a tool to investigate how network organization and dynamics jointly support learning and functional specialization.

Here we combine connectomics and reservoir computing to investigate the link between network organization, dynamics and computational properties in human brain networks. We construct neuromorphic artificial neural networks endowed with biologically realistic connection patterns derived from diffusion weighted imaging. We then train these connectome-informed reservoirs to perform a temporal memory task. To evaluate how memory capacity depends on both network structure and dynamics, we parametrically tune the dynamical state of the network, driving the network to transition between stable, critical and chaotic dynamics. We assess memory capacity of empirically-derived connectomes against two null models, and show that the underlying topology and modular structure of the brain enhance memory capacity in the context of critical dynamics. Throughout the report, we focus on the role of *intrinsic networks* [12, 32, 106, 115, 125, 141]: large-scale functional systems of the brain thought to be the putative building blocks of higher cognition [14, 15, 145]. Finally, we systematically investigate how memory capacity depends on the interaction between specific network attributes and dynamics.

## RESULTS

### Learning in neuromorphic networks

Reservoir computing was initially proposed as a computational framework to model how cortical circuits extract information from the spatiotemporal patterns of neural activity elicited by continuous streams of external stimuli [24]. Nowadays reservoir computing is primarily used as an artificial recurrent neural network architecture in its own right [80]. In its simplest form, the “vanilla” reservoir computing architecture consists of a recurrent network of interacting nonlinear neurons, known as the *reservoir*, and a linear readout module. In the present report, we use empirically-derived patterns of human brain connectivity to constrain the connections within the reservoir. Specifically, we apply deterministic streamline tractography on diffusion spectrum MRI data (*n* = 66 subjects. Data source: https://doi.org/10.5281/zenodo.2872624) to reconstruct individual high-resolution human connectomes (1015 nodes; 1000 cortical and 15 subcortical regions). To reduce the impact of false positives and false negatives, as well as the effect of inconsistencies in the reconstruction of subject-level connectomes on network measures, we generate group-level consensus networks that retain the topological characteristics of individual subject networks [19, 33, 110].

We use the reservoir framework to quantify the capacity of individual brain regions to encode temporal information during a memory task (Fig. 1a). In this task a uniformly distributed temporal signal is introduced into the reservoir through a set of *input nodes* (blue). The signal propagates through the network, giving rise to complex patterns of neural activity across the reservoir. The activity time-series of a set of *readout nodes* (purple) within the reservoir are then used to train a readout linear unit to reproduce a delayed version of the input signal. Performance in this task thus depends on the ability of the readout nodes to encode the temporal properties of the external stimuli. In other words, performance depends on the ability of readout nodes to represent past and present inputs in their current activation state, which we refer to as *memory capacity* (see *Methods* for details).

**Figure 1.**
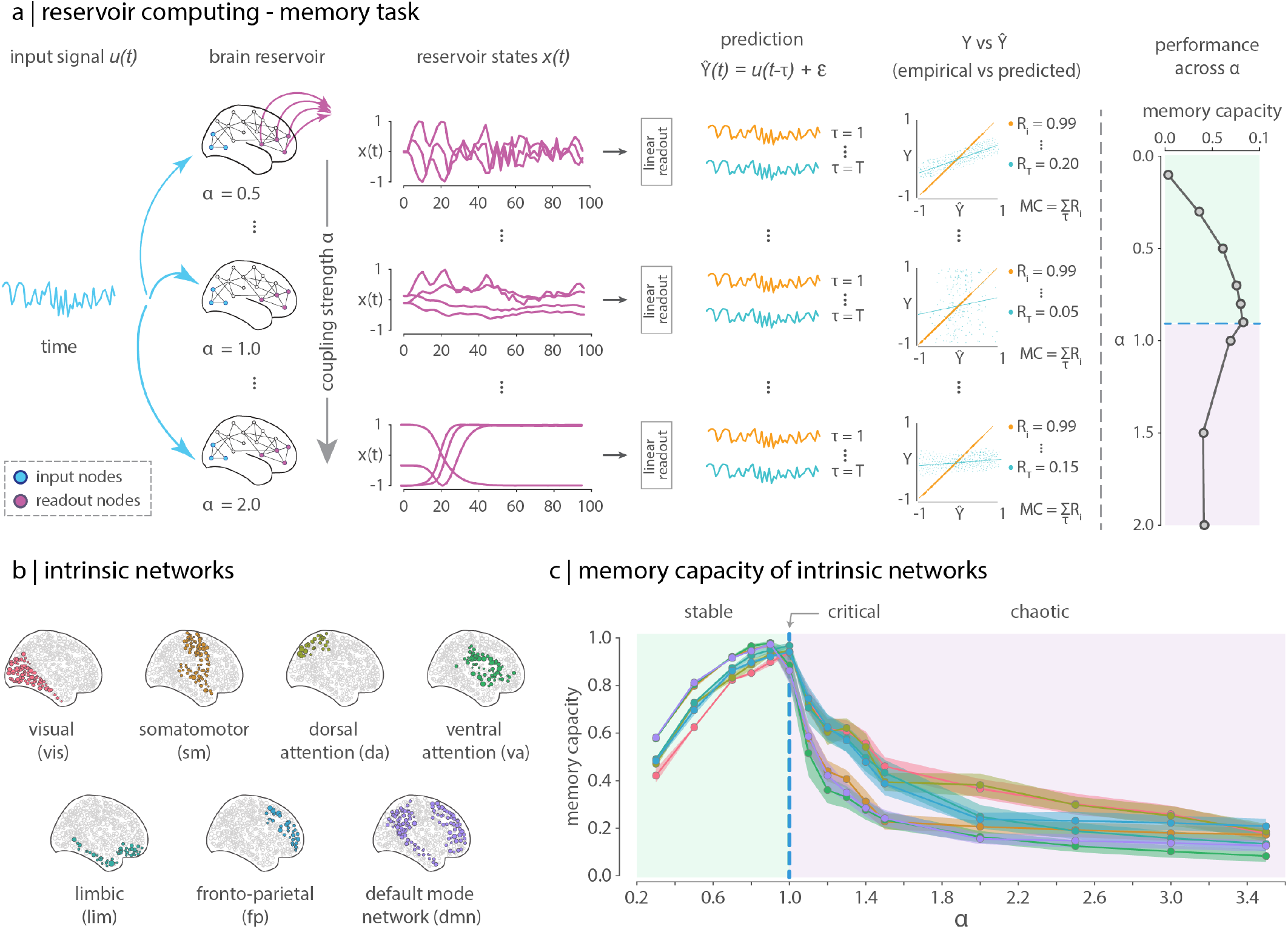
Measuring the memory capacity of biological neural networks. By using biologically-inspired connectivity measures obtained from MRI connectomics data, we leverage reservoir computing models to investigate the effect of network architecture on the encoding capacity of large-scale brain networks in a memory task. (a) In a memory task, a uniform random input signal (*u*(*t*)) is introduced to the reservoir through a set of input nodes (blue nodes). The input signal propagates through the network, activating the states of the units within the reservoir. The activation states (reservoir states *x*(*t*)) of the readout nodes (purple nodes) are then retrieved from the reservoir, and are used to train a linear model to reproduce a delayed version of the input signal at different time lags (*Y* = *u*(*t − τ*)). Memory capacity (*MC*) is estimated as the sum across time lags (*τ*) of the absolute correlation (*R*) between the predicted 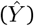 and the time-lagged target signal (*Y*). The dynamics of the reservoir are controlled with *α*, a parameter that scales the spectral radius (largest eigenvalue) of the connectivity matrix, while uniformly modifying its coupling strength. A memory capacity estimate is obtained for every value of *α*. (b) Lateral projection (right hemisphere) of the Yeo-Krienen intrinsic network partition [141]. The procedure described in (a) was independently applied to each of the seven intrinsic networks. (c) Distribution (across bootstrapped consensus matrices; see *Methods* for details) of the memory capacity as a function of *α* for each of the seven intrinsic networks.

We selected readout nodes using intrinsic networks, [12, 32, 106, 125, 141] (Fig. 1b). Intrinsic networks are a connectivity-based partition of the brain into groups of areas with coherent time courses and similar function [159]. As such, they are thought to represent the putative building blocks of higher cognition, and provide a convenient and meaningful way to divide the brain into large-scale systems. In the present study, we measured the memory capacity of the brain across seven intrinsic networks reported by Yeo, Krienen and colleagues [141]. Subcortical regions were set as input nodes for all experiments (results using alternative input nodes are shown in *Sensitivity Analysis*). To provide confidence intervals for the memory capacity estimates of intrinsic networks, we generate 1000 group-consensus matrices by bootstrap resampling individual subjects (see *Methods* for details).

The computational properties of the reservoir depend on both its architecture and its dynamics [25, 87]. Here, the reservoir consists of a recurrent neural network of discrete-time, nonlinear threshold units. We parametrically tune the dynamics of the reservoir by scaling the largest eigenvalue of the connectivity matrix (parameter *α*; see *Methods*). The dynamics of the reservoir are considered to be stable when *α <* 1, and chaotic when *α >* 1. When *α* ≈ 1 the dynamics are said to be critical or at the edge of chaos [118]. Fig. 1c shows the mean (solid lines) memory capacity and the standard deviation (shaded regions) across bootstrapped samples for each of the seven intrinsic networks as a function of *α*. As expected, the memory capacity is greater when the dynamics of the reservoir are stable (*α <* 1), and decreases as the dynamics evolve towards a chaotic state (*α >* 1) [118]. Maximal memory capacity is attained when the dynamics of the reservoir are at the transition between the stable and chaotic regimes (*α* ≈ 1). This general behaviour is observed across all the intrinsic networks, with more prominent differences between them in the chaotic regime. The greater within-system variability in memory capacity at *α >* 1 is consistent with the notion that the system is in a chaotic state, while capacity in the stable regime is less variable.

### Memory capacity of the human connectome

We first assess how memory capacity depends on connectome architecture and dynamics. We averaged the memory capacity across functional systems to provide an overall estimate for the memory capacity of the brain. Fig. 2a shows the distribution of mean memory capacity across functional systems as a function of *α*. Consistent with the results from the previous section, we find the greatest performance close to the edge of chaos, with memory capacity generally > 0.8 at *α* values close to 1. We next ask how performance depends on network topology and on the modular organization captured by the intrinsic networks.

**Figure 2.**
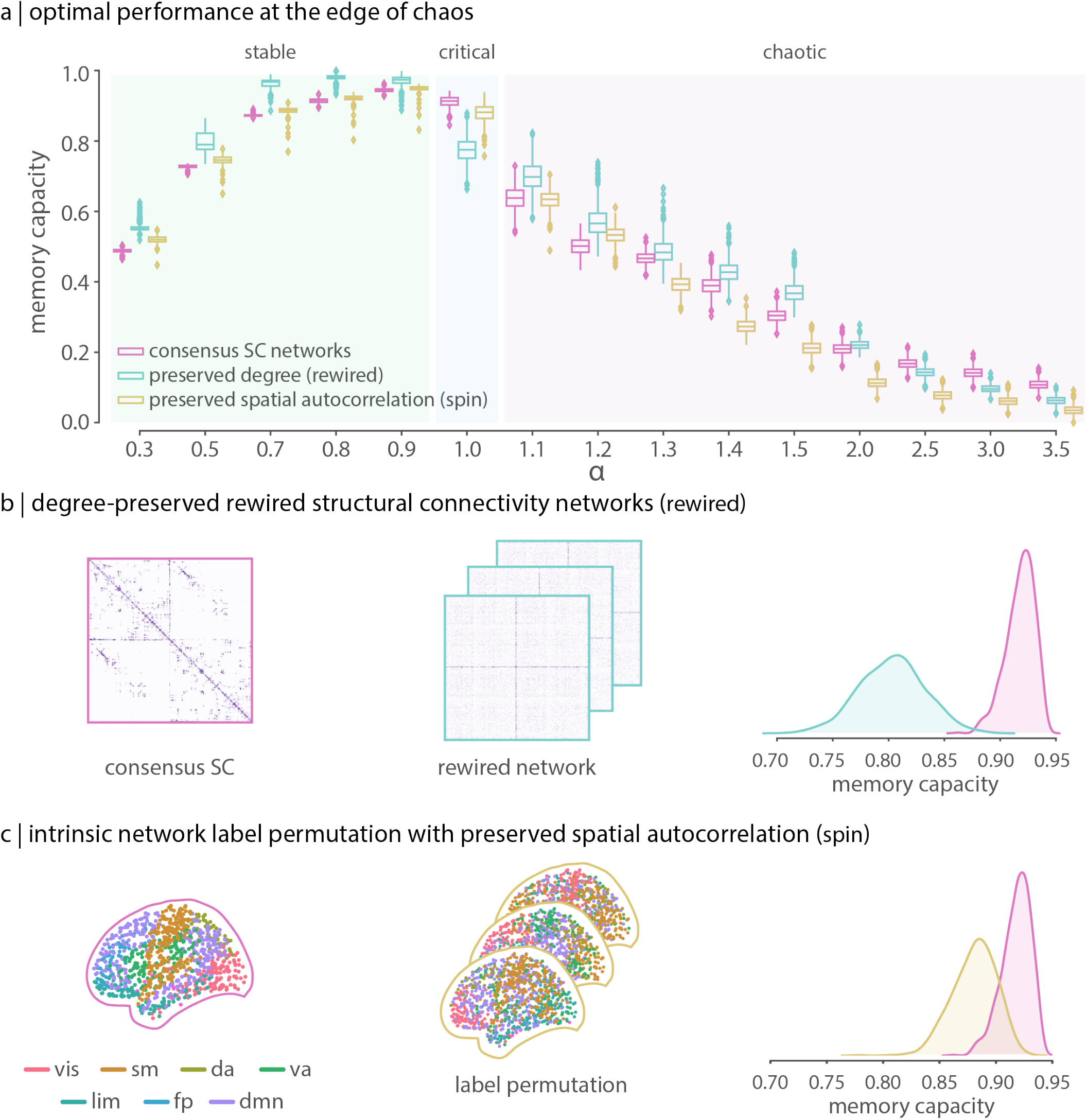
The unique features of the connectome drive memory capacity in the critical state. (a) Distribution (across bootstrapped consensus matrices) of the memory capacity of the human connectome (magenta) as a function of *α*. Whole-brain-level memory capacity estimates were calculated as the average memory capacity across intrinsic networks. At the *edge of chaos* (*α* = 1), memory capacity of the human connectome is optimal compared to two alternative null models ((b) and (c); cyan and yellow, respectively). (b) To determine whether memory capacity estimates depend on the underlying connectivity structure, a null distribution is constructed by randomly rewiring pairs of edges, while preserving network size, density, degree sequence and node-level intrinsic-network assignment (cyan in panel (a); referred throughout the text as *rewired*) [86]. (c) To evaluate the extent to which the partition of the connectome into seven intrinsic networks is relevant for the task at hand, a null distribution is constructed by spherical projection and random rotation of the intrinsic network labels, preserving their spatial embedding and autocorrelation (yellow in panel (a); referred throughout the text as *spin*) [1, 85].

To assess the extent to which memory capacity depends on the underlying network topology, we constructed a population of null networks by randomly rewiring pairs of edges in the structural network, preserving the density, degree sequence and the intrinsic network assignment of the nodes ([86]; Fig. 2b). Using this null model, we performed the same memory task to measure the memory capacity of each of the seven functional systems, and then averaged across them to provide a network-level memory capacity estimate. This procedure was applied on 1,000 rewired networks to construct a distribution of the memory capacity of the network under the null hypothesis that network memory capacity is independent of the underlying connectivity patterns. We then compare brain networks against the rewired models using the Wilcoxon-Mann-Whitney two-sample rank-sum test. We find that empirical brain networks have significantly better performance than rewired networks at criticality (medians in empirical and rewired networks are 0.92 and 0.80, respectively, *P_rewired_ <* 10^−4^ two-tailed, and effect size = 99%; Fig. 2a); at other dynamical regimes (stable and chaotic), the connectome generally has lower capacity. Altogether, these results suggest that connectome topology optimally functions as a reservoir when dynamics are critical or at the edge of chaos. We next sought to assess whether the partition of the connectome into seven intrinsic networks is functionally meaningful. To do so, we used a spatially-constrained label-permutation null model that randomly permutes intrinsic network labels while preserving their spatial autocorrelation, testing the hypothesis that the memory capacity of the brain depends on its functionally-defined modular structure, and not on trivial differences in size, coverage or symmetry of intrinsic networks ([1, 85]; Fig. 2c). Again, we use the Wilcoxon-Mann-Whitney two-sample rank-sum test to compare the empirical intrinsic network partition against the spatially-constrained permutation model at the edge of chaos. We find that the partition provided by empirical intrinsic networks significantly outperforms the spatially-constrained label-permutation model at criticality (medians in the empirical and permuted partitions are 0.92 and 0.89, respectively, *P_spin_ <* 10^−4^ two-tailed, and effect size = 88%; Fig. 2a), as well as in the chaotic regime. This suggests that the modular organization of the brain in functional systems defined by intrinsic networks constitutes a computationally-relevant feature of the human connectome [14, 15].

### Connectomes optimize computation-cost trade-offs

Previously, we observed that the human connectome outperforms the rewired model only when the dynamics are critical. In the stable and chaotic regimes, randomly rewired networks outperform the empiricallly-derived architecture (Fig.2a). However, this null model does not account for the fact that the brain is a spatially-embedded network, with finite metabolic and material resources [23, 132]. As a result brain networks are characterized by a prevalence of shorter, low-cost connections [57, 109]; by contrast the random flipping of edges in rewired networks skews the connection length distribution towards longer connections resulting in networks with greater wiring cost than empirical brain networks (Fig.3a) [18, 20]. Fig.3b shows memory capacity of real and rewired networks, normalized by wiring cost. When we account for the cost of wiring, empirical connectomes outperform rewired networks across the three dynamical regimes, suggesting that brain network topology optimizes the trade-off between computational capacity and cost.

**Figure 3.**
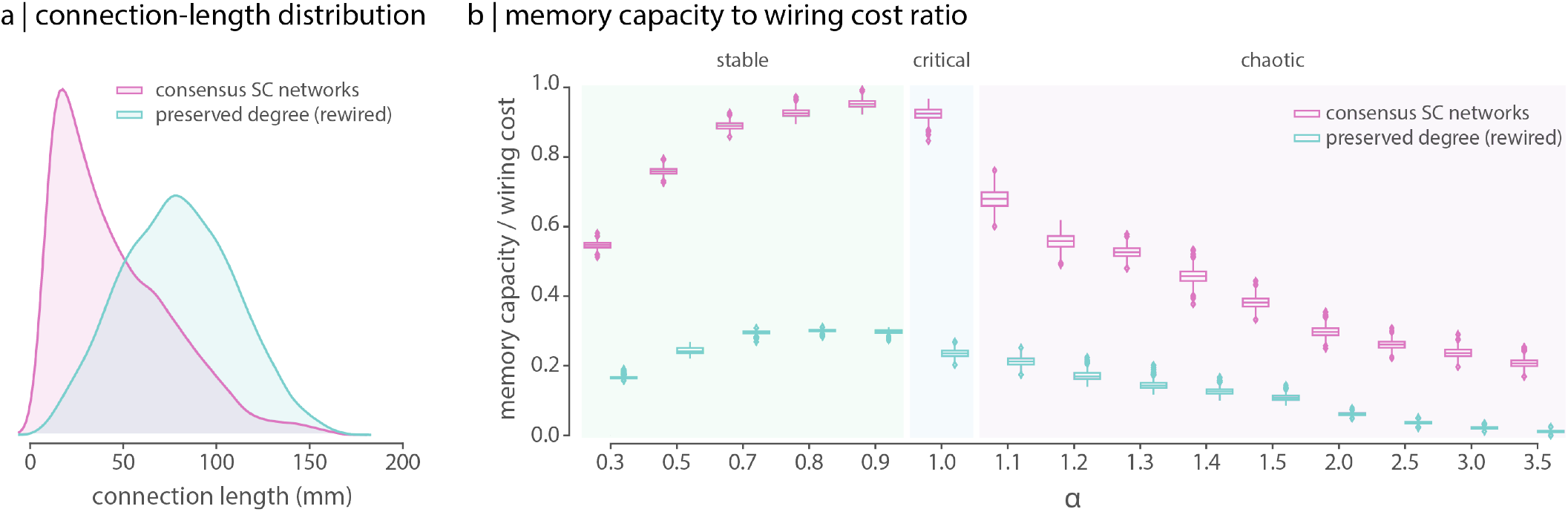
Accounting for the cost of wiring. (a) Connection length distribution for a sample consensus network (magenta) compared to a sample rewired network (cyan). Euclidean distance between brain regions was used as a proxy for connection length. (b) Memory capacity to wiring cost ratio. The wiring cost was estimated as the sum of the connection lengths weighted by the connectivity values, which are proportional to the number of streamlines between brain regions. When the wiring cost is taken into account, the empirical network outperforms the rewired null model across all dynamical regimes.

### Memory capacity of intrinsic networks

We next investigate the memory capacity of individual functional systems. Fig. 4a shows the distribution of memory capacity of individual intrinsic networks, as well as the corresponding null distributions generated by rewiring and spatially-constrained label permutation. Performance is stratified according to dynamical regime, from stable (mean performance for *α* = [0.3, 0.5, 0.7, 0.8, 0.9]) to critical (performance for *α* = 1.0) to chaotic (mean performance for *α* = [1.5, 2.0, 2.5, 3.0, 3.5]). To establish a clearer distinction between the critical and the chaotic regimes, we selected *α* values above 1.5 to characterize chaotic dynamics (results for intermediate values of *α* in the range 1.1 to 1.5 are provided in Fig.S1). Throughout subsequent analyses, dynamical regimes are defined using the same ranges across *α* values provided here.

**Figure 4.**
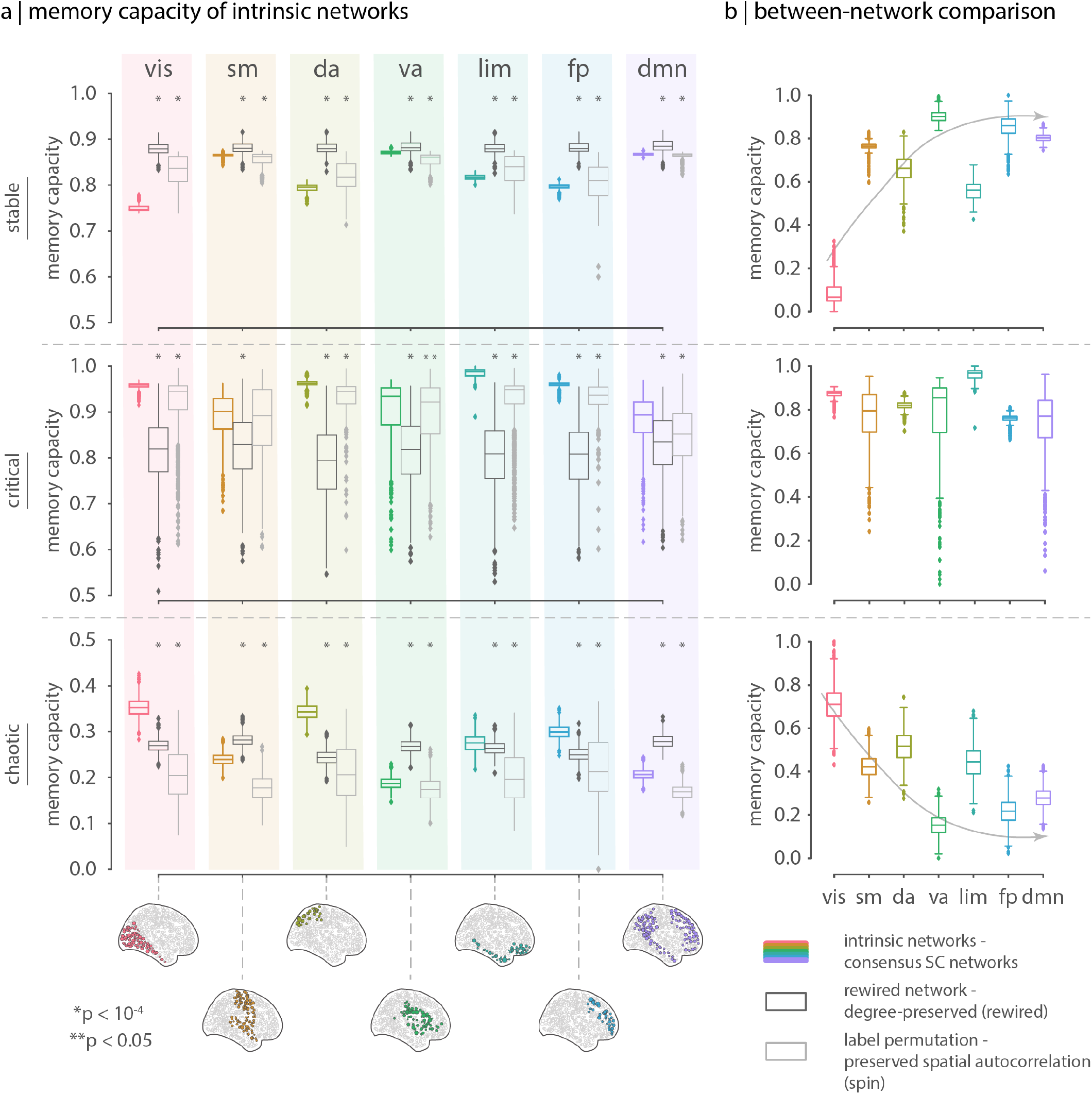
Generalized optimal memory capacity across intrinsic networks. (a) At the *edge of chaos*, memory capacity estimates of the empirical network are consistently and significantly higher and optimal across functional systems, compared to their analogue estimates computed on the rewired and spatially-constrained null models. In other words, optimal performance of the human connectome in the critical state is not driven by a few networks, but is rather generalized across the brain. (b) Memory capacity in recurrent neural circuits largely depends on the size of the network and the amount of positive feedback, which are in this case proportional to the number of nodes and the number of internal connections, respectively [56]. A direct and fair between-network comparison would require removing the effect of these factors from the memory capacity estimates of individual networks. Using a linear model, the effect of relative connection density was regressed out (see S2 for further details). To facilitate the comparison, memory capacity estimates were scaled between 0 and 1 using the minimum and maximum values within each dynamical regime. When compared to each other, memory capacity is in general quite homogeneous across functional systems in the critical regime, but it is highest for the limbic system, maybe reflecting an anatomically-mediated predisposition for memory encoding [11, 97, 98, 116, 160]. Interestingly, in the stable and chaotic regimes, memory capacity estimates differentiate somatosensory from higher association areas [58, 84, 92, 93].

We use the Wilcoxon–Mann–Whitney two-sample rank-sum test to compare the performance of empirical intrinsic networks against their analogues in the rewired and label-permutation null models. Consistent with the intuition gleaned from the global performance shown in the previous section, we find significantly greater memory capacity for most functional systems in the critical regime, at the edge of chaos (mean memory capacity > 0.9 for all networks), but poorer or inconsistently better performance in the stable and chaotic regimes (across dynamical regimes, *P_rewired_ <* 0.05 and *P_spin_ <* 0.05 two-tailed, compared to the rewired and the label-permutation models, respectively).

How do functional systems compare with each other in terms of memory capacity? Making direct comparisons between intrinsic networks is challenging, as memory capacity in individual networks will be partially driven by features such as network size (i.e., number of nodes) and number of internal connections (S2) [56, 167]. To account for differences among networks due to density, we normalize their memory capacity by their relative connection density (see S2).

Fig. 4b shows the density-normalized network-specific memory capacity at each of the three dynamical regimes. In the critical regime, we observe few differences among networks, with the greatest memory capacity in the limbic network, perhaps reflecting an anatomically-mediated predisposition for memory encoding [11, 97, 98, 116, 160]. In the stable and chaotic regimes we observe diametrically opposite orderings, such that networks with greater capacity in the stable regime have lower capacity in the chaotic regime, and vice versa. This suggests an interplay between network topology, dynamics and memory capacity, a phenomenon we study in greater detail in a subsequent section. Interestingly, the axis or trend line along which the memory capacity of intrinsic networks fluctuates in these regimes broadly resembles the putative unimodal-transmodal hierarchy [58, 84, 92, 93], differentiating sensory networks (visual and somatomotor) from association networks (default and frontoparietal), suggesting a potential link between the hierarchical organization of the cortex and cognitive capacity [139, 164].

### Information transfer across the brain

The performance of intrinsic networks in the memory task provides a measure of how the macroscopic human connectome supports temporal information storage. This metric, however, tells us little about how information changes as it travels from one region to the other in the brain. We thus next investigate how temporal information content (i.e., memory capacity) is transformed as signals propagate through the connectome during learning. Specifically, we compare the memory capacity of each intrinsic network in an *encoding* and a *decoding* experimental set up (Fig. 5a). For encoding, we measure the memory capacity of the information *sent* by the network to the rest of the brain, that is, the signals generated by the network itself. For decoding, we measure the memory capacity of the information *received* by the network, that is, the signals generated by its directly connected neighbours. We call these metrics the encoding and decoding capacity, respectively, of an intrinsic network (see *Methods* for interpretation, rationale and further considerations on these metrics).

**Figure 5.**
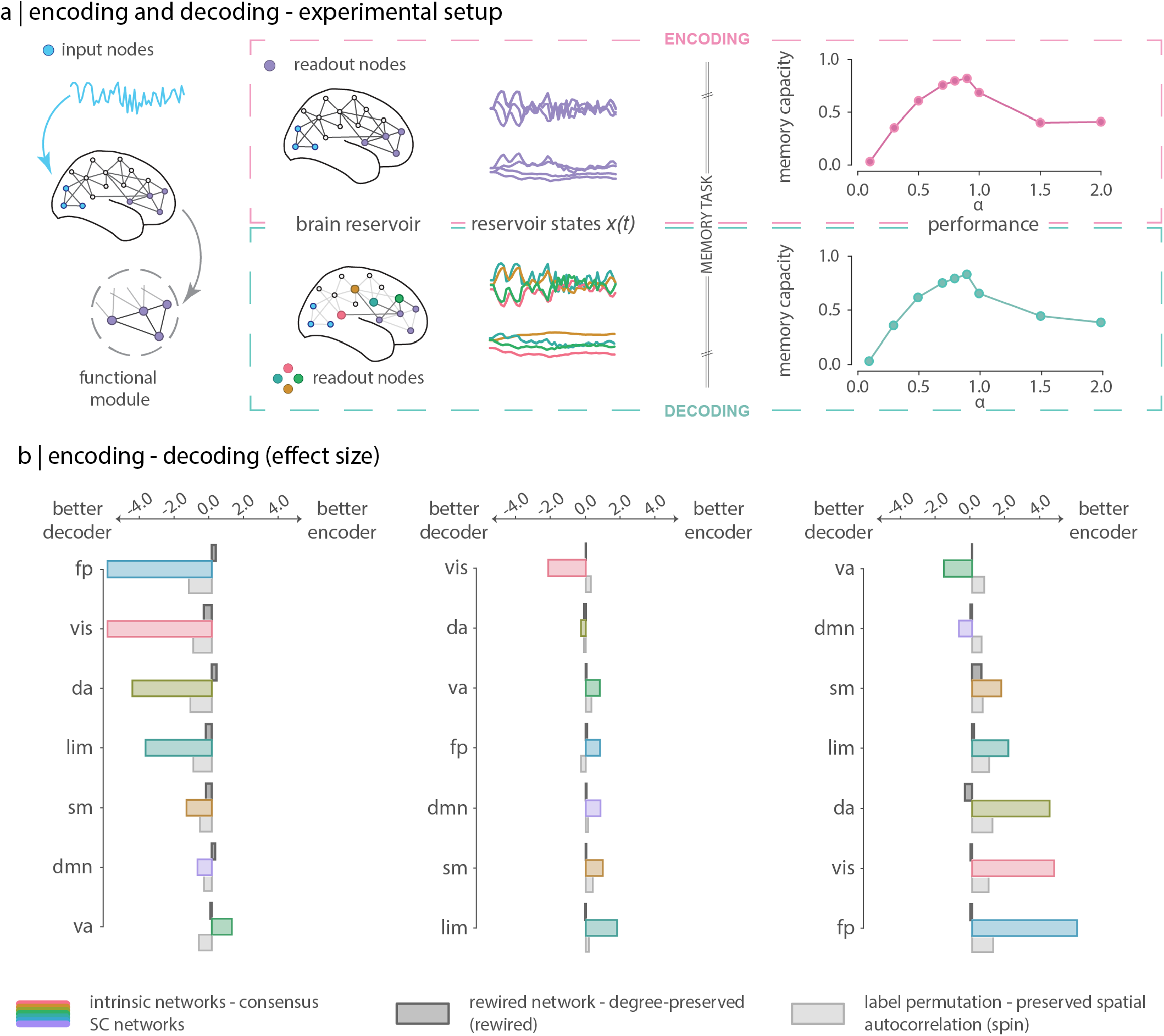
Encoding and decoding capacity of functional systems. The difference between the encoding and decoding capacity of a network quantifies the extent to which information is transformed as it propagates through the network. (a) The encoding (decoding) capacity of a network is equivalent to its memory capacity when it operates as an encoder (decoder). The coding role of the network is determined by the selection of the readout nodes. In the encoding setup (top panel) readout nodes are the nodes that belong to the network; they are a proxy for the information *sent* by the network. In the decoding setup (bottom panel) readout notes correspond to the direct neighbours of the network; they are a proxy for the information *received* by the network. (b) Bar plots show the effect size of the difference between encoding and decoding, quantified as the *Cohen’s D* estimator consistent with a 1-sample *t*-test. Compared to the rewired and spatially-constrained null models, the human connectome presents overall a higher encoding - decoding asymmetry.

Differences between encoding and decoding capacity quantify the extent to which temporal information content is transformed as signals travel through a functional network [122]. The key finding is that connectome topology supports functional specialization by promoting regional heterogeneity of temporal information content. To determine whether encoding and decoding are statistically different from one another, we conduct a two-tailed, one-sample *t*-test under the null hypothesis that the mean of the difference between encoding and decoding capacity is equal to zero (*H*_0_: *μ_enc.−dec._* = 0). In favor of regionally heterogeneous dynamics, we find that, across all dynamical regimes, the encoding - decoding differences of intrinsic networks are significantly different from zero (*P <* 10^−4^, Bonferroni corrected). Fig. 5b shows the effect size of the difference between the encoding and decoding capacity of intrinsic networks across dynamical regimes. We quantify effect size as the Cohen’s D estimator consistent with a one-sample *t*-test for means (*H*_0_: *μ_enc. − dec._* = 0). We use this measure to compare the empirical network against the rewired and label-permutation models. Importantly, across all functional systems and dynamical regimes, we observe significantly greater asymmetry between encoding and decoding for the empirical connectome compared to the rewired and spatially-constrained null models. This suggests that both the brain’s network connectivity, and its modular organization into large-scale systems, play a role in optimizing how information is transformed as signals propagate through the connectome, thus promoting regional heterogenity and supporting functional specialization.

At criticality – when overall connectome performance is optimal – we observe smaller but still significant differences between encoding and decoding. The reduction in the difference between encoding and decoding capacity at the critical point suggests a homogenization of information content across the connectome (integration). At the same time, the magnitude of the difference between encoding and decoding is significant (minimum absolute effect size > 0.25), indicating a heterogeneous distribution of information across functional networks (segregation). In other words, at the critical point, there exists a balance between integration and segregation of the information across the connectome [127, 128]. Interestingly, the functional system with the greatest difference between encoding and decoding capacity at criticality is the limbic system, again suggesting an anatomical basis for the well-studied involvement of this system in memory function [11, 97, 98, 116, 160].

Finally, it is noteworthy that most functional systems are computationally flexible, performing better as decoders in the stable regime (*decoding > encoding*), and better as encoders in the chaotic regime (*decoding > encoding*). This is the case for all intrinsic networks (visual, somatomotor, dorsal attention, limbic and frontoparietal; Fig. 5b, top panel), except the ventral attention and the default mode network systems. Interestingly, the default mode network is the only functional system that acts as a decoder in both the stable and the chaotic regimes, while its encoding capacity is expressed only at criticality, consistent with contemporary theories that this functional system supports global monitoring and information integration [40]. Overall, results suggest that topology and dynamics interact to support switching of the direction in which temporal information content accumulates in functional networks, endowing these systems with computational flexibility.

### Interplay between network topology, dynamics and memory capacity

A recurring theme in the previous sections was that memory capacity depends on the interaction between network topology and dynamics. Here we relate memory capacity to multiple (local- and global-level) attributes of the connectome graph across the three dynamical regimes. Fig. 6a shows the (distribution of) correlations between memory capacity and local graph attributes; attributes are first computed for individual nodes and then averaged within intrinsic networks. Fig. 6b shows the correlations between memory capacity and global graph attributes. All graph properties were computed using the weighted structural connectivity network. Briefly, graph attributes include measures of “connectedness (strength), clustering, centrality (betweenness), connection diversity (participation), global efficiency (characteristic path length), and modularity (see *Methods* for definitions).

**Figure 6.**
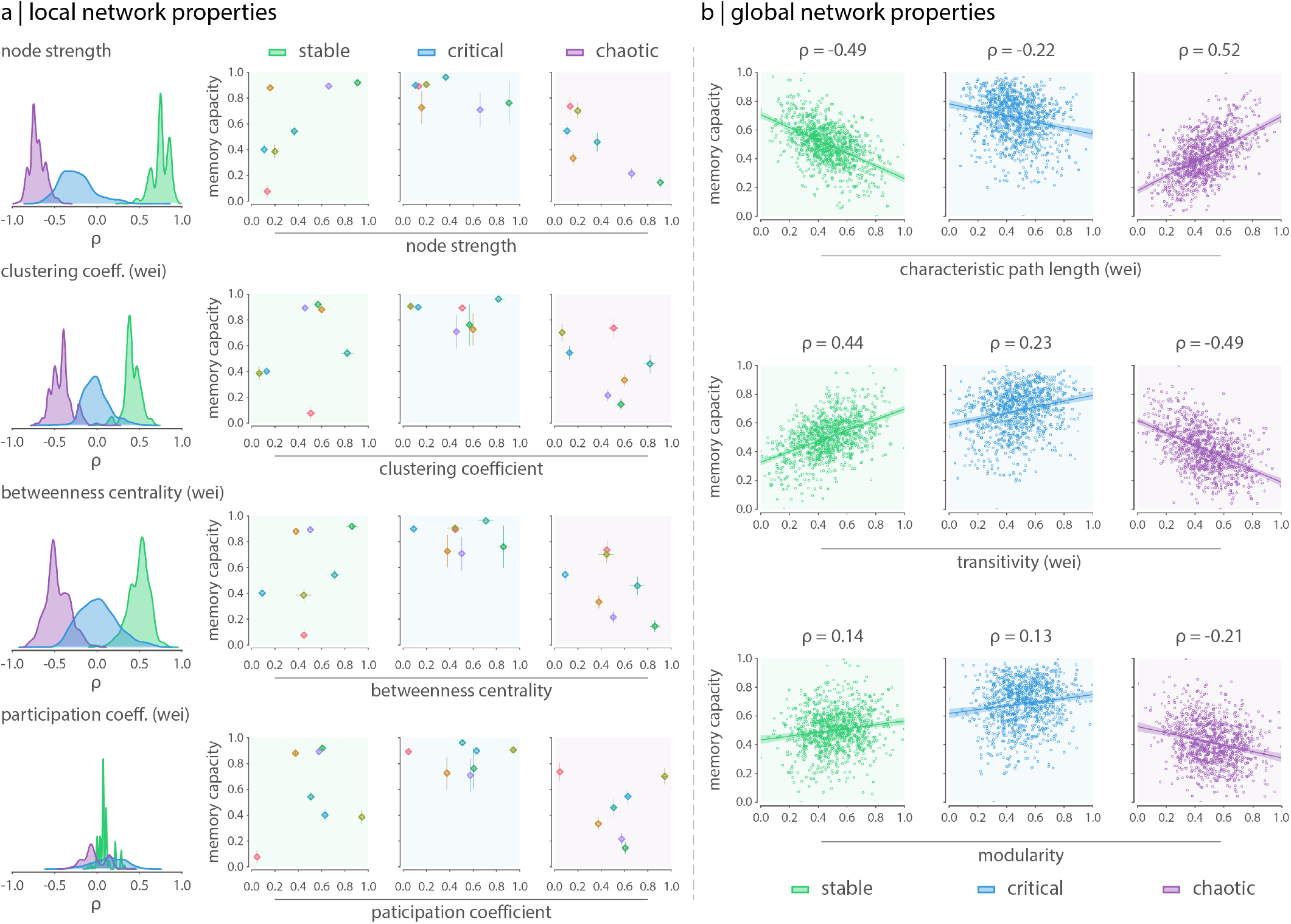
Contribution of network topology to memory capacity is a function of the dynamics. Local and global topological attributes interact with network dynamics to shape the memory capacity of the human connectome. (a) Correlation between local network topology and the memory capacity of intrinsic networks across dynamical regimes. Local features, including node strength, clustering coefficient, betweenness centrality and participation coefficient, were estimated at the node level and then averaged within each intrinsic network. Left column shows the distribution (across bootstrapped samples) of the Spearman correlation between average local network properties and the memory capacity of intrinsic networks. Right column shows the scatter plots of memory capacity vs network attributes; markers (error bars) represent the average (standard deviation) across bootstrapped samples. (b) Correlation between global network topology and memory capacity of the human connectome across dynamical regimes. Global features include characteristic path length, transitivity and modularity.

We note three key results. First, we observe moderate to high correlations between graph attributes and memory capacity, suggesting that network structure influences performance. Second, the direction of that influence depends on the dynamics, with correlation signs switching from one regime to another. In other words, structure does not simply drive dynamics, and thus performance. Rather, structure and dynamics interact to drive performance. Third, at criticality, memory capacity is much less dependent on the topology, compared to the stable and chaotic regimes, where the correlations are typically greater. That is, at criticality, memory capacity transcends topological features and becomes more dependent on global network dynamics.

### Sensitivity analysis

Throughout our analyses, subcortical regions, which include structures such as the thalamus, the basal ganglia, the hippocampus, the amygdala and the brain stem, served as input nodes in all experimental set ups. Nevertheless, not all these regions actually constitute biologically-plausible relay stations for incoming sensory signals. From all the subcortical structures, the thalamus is known to play an important role in relaying sensory and motor signals from the rest of the body to the cerebral cortex and to other cortical structures [120]. Therefore, we repeated our analyses under a more realistic experimental set up that considers only those subcortical regions comprising the left and right hemisphere sections of the thalamus. Results are shown in Fig. S3 and S4. Although memory capacity estimates change from one type of experimental set up to the other, the human brain network still displays maximum and optimal memory capacity in the critical state compared to the rewired (*P_rewired_* = 0.0 and effect size = 99% two-tailed, Wilcoxon-Mann-Whitney two-sample rank-sum test) and label-permutation (*P_spin_* = 0.0 and effect size = 99% two-tailed, Wilcoxon-Mann-Whitney two-sample rank-sum test) models. This is true for the whole brain, and also for individual intrinsic networks. In fact, under this more realistic scenario, statistical significance and effect sizes are even stronger.

Likewise, all analyses presented thus far were conducted on a particular parcellation of brain regions. We repeated our analyses at a lower resolution parcellation (448 cortical and 15 subcortical regions) to assess the extent to which our results depend on network scale. Results are shown in Fig. S5 and S6. At this lower scale, the overall memory capacity of the brain in the critical state is still statistically higher compared to the rewired and label-permutation models. However, this is not the case for all intrinsic networks. The ventral attention, limbic and default mode network systems display statistically higher performance compared to the null models. The visual and somatomotor systems present a statistically higher performance compared to the rewired model, but significantly lower performance compared to the label permutation model. Finally, both the dorsal attention and fronto-parietal networks present statistically lower performance compared to the rewired null model, but only the fronto-parietal system presents a significantly higher performance compared to the label permutation model. Memory capacity is a property that is strongly tied to the underlying within-network connectivity and relative density of each intrinsic network. Because connectivity details involving these two factors can be easily dismissed when the networks are reconstructed at a lower resolution parcellation (since each brain region occupies now a larger volume), subtle differences found at the intrinsic network level are not surprising.

## DISCUSSION

In the present report we used reservoir computing to study how network structure and dynamics shape learning and computations in networks with architectures based on the human connectome. Using neuromorphic neural networks, we show that empirically-derived architectures perform optimally at criticality, and excel at balancing the trade-off between adaptive value and cost, regardless of the dynamical regime. We find that performance is driven by network topology, and that the modular organization of large-scale functional systems is computationally relevant. Throughout, we observe a prominent interaction between network structure and dynamics, such that the same underlying architecture can support a wide range of learning capacities across dynamical regimes.

By studying artificial neural networks with connectome-based architectures, we begin to reveal the functional consequences of brain network topology. Numerous studies point to a unique set of organizational features of brain networks [7, 112, 130, 146], but how these features influence the neural computations that support cognition remains unknown. By training networks with biologically-realistic connectivity to perform a simple memory task, we show that connectomes achieve superior performance compared to populations of networks with identical low-level features (density, degree sequence) but randomized high-level topology.

The present work highlights a symbiotic relationship between structure and dynamics. Namely, the unique topological features of the brain support optimal performance only when dynamics are critical, but not when they are stable or chaotic. Multiple accounts posit that the brain, akin to many other naturally occurring complex systems, operates at the interface of order and disorder. In this critical regime, at the “edge of chaos”, dynamic transitions are optimized, and thought to confer adaptive benefits, including greater dynamic range and selective enhacement of weak inputs [50, 121, 150]. In addition, multiple empirical studies have demonstrated that neural dynamics exhibit fluctuations consistent with a networked system at criticality [28, 35, 69, 137].

In the context of reservoir computing, a link between the computational power of the reservoir and the edge of chaos has been established in multiple theoretical studies [16, 72, 74, 75]. The reservoir has the potential to work as a universal function approximator as long as it can hold (memory) and nonlinearly transform (separation) information about past external stimuli in its high-dimensional transient states [81]. The balance between these two properties, namely the *fading memory property*, which is enhanced by stable dynamics, and the *separation property*, which is supported by chaotic dynamics, is optimal at the edge of chaos. At this critical point, where the transition between stable and chaotic dynamics occurs, the perfect balance between order and disorder delivers the optimal trade-off between memory and separation. The fact that brain networks perform better at criticality (compared to the rewired network model) suggests that network topology may be configured to negotiate that optimal trade-off between these two computational properties, and therefore maximize computational capacity.

Computational power and information processing, however, might not be the only features that brain networks seem to optimize. The brain is a spatially embedded network with finite metabolic and material resources, and as such, there are physical demands that biological networks should satisfy [23]. For instance, the growth and maintenance of axonal projections entails material and energetic costs. These wiring costs increase with the length of inter-neuronal connections and manifest as a prevalence of short-range connections [57, 109]. Therefore, studies of the computational, topological and dynamical features of the brain must also consider the cost imposed by its geometrical constraints [17, 20, 65]. A salient finding in the present report is that, when wiring cost is taken into account, brain networks outperform randomized networks in all dynamical regimes. The fact that empirical networks achieve higher computational power per unit cost compared to the rewired networks supports the idea that brain networks are highly economical: they maximize adaptive value while minimizing wiring cost [9].

From an engineering perspective, these results present a new direction for designing neuromorphic or biomimetic artifical neural networks. Deep artificial neural networks have myriad applications in modern machine learning and artificial intelligence. Despite superficial similarity with macro-level connectivity in the brain, architectural design of deep networks is typically ad hoc and problem-specific. Moreover, there are increasing efforts in both academia and industry to develop next-generation neuromorphic hardware devices and chips. One such effort is the construction of physical *reservoirs* [138]. These include reservoirs built from analog circuits [6, 77, 126, 166], field-programmable gate arrays (FPGA; [1, 2, 4, 5, 156]), very large-scale integration circuits (VLSI; [103, 105, 111]), memristive networks [13, 41, 68, 71, 123, 134, 161], and photonic or opto-electronic devices [66, 67, 73, 148, 149, 165]. The architectures of these algorithms and systems, however, rarely take advantage of emerging understanding of connection patterns in biological networks. We envision that the present work serves as a foundation for future efforts to design biologically-inspired artificial neural networks and physical neuromorphic hardware. Due to their physical nature, neuromorphic systems are conditioned to similar material and energetic costs as biological neural networks. Because of this, the cost-effective design of these information processing systems could largely benefit from insights gained about the economical principles of brain network organization [9]. Ultimately the present paradigm could be used to systematically map combinations of network attributes and dynamical regimes to a range of computational functions, and may ultimately help to identify design principles for engineering better networks and circuits.

More broadly, the present work addresses the question of the structure-function relationship in the brain, but with a focus on computation. Over the past 20 years, multiple forward models have been proposed to link structural connection patterns to functional connection patterns [135], including statistical models [91, 94], communication models [30, 43, 45, 95] and neural mass models [22, 34, 54, 114]. These models have been successful in predicting an array of empirical phenomena, including static and dynamic functional connectivity [55]. However, these models are largely phenomenological, and focused on predicting an emergent property of the system (intrinsic functional connectivity), rather than explaining how the system computes and approximates functions in the external environment. The present work represents a step towards mapping network structure more directly to its computational function.

In particular, we provide evidence that the intrinsic networks of the brain constitute computationally relevant systems. Despite their widespread use as a frame of reference in cognitive neuroscience, their precise cognitive function is not completely understood. Although they are typically labeled based on prior domain knowledge from functional neuroanatomy [145], these networks are primarily defined as groups of neuronal populations with coherent time courses. Multiple studies have reported correlations between the expression of these networks and individual differences in behaviour [89], but how they contribute to computation in the brain is unknown. The present results demonstrate that imposing this intrinsic functional network partition on the structural connectome yields better performance, implying that these networks are indeed functionally meaningful [15, 49]. As one salient example we find that, at criticality, the limbic network consistently presents the greatest memory capacity, and the greatest difference between encoding and decoding, suggesting that the contribution of this network to memory function naturally arises from its embedding in the macroscale anatomical architecture of the brain [11, 97, 98, 116, 160]. We also observe that the interaction between topology and dynamics shapes how information accumulates in the network. Interestingly, the direction in which information accumulates in non-critical regimes mirrors the characteristic ordering of functional systems along the unimodal-transmodal hierarchy [58, 84, 92, 93], suggesting a possible mechanism to flexibly switch between bottom-up and top-down information flow.

The present paradigm can be extended to investigate a wider range of cortical functions to further our understanding on how functional specialization emerges from network structure. By training brain-isnpired network architectures to implement other types of tasks, we can probe other modes or classes of computation. For example, temporal-sequence learning tasks, mimicking speech recognition, depend on the network’s ability to occupy multiple states and match these states to the signal inputs, thus accessing a different computational property. We thus envision future efforts to link architecture and computation in brain networks, analogous to modern meta-analytic methods that map regional activation to cognitive tasks [39, 42, 104, 162]. Mapping network structure and dynamics to fundamental blocks of computation may allow us to build a comprehensive structural-functional ontology, relating network structure to computational properties and, ultimately, to cognitive function.

We finally note that the present results should be interpreted with respect to several limitations. First, to focus on the question of how network structure influences learning, we implemented a model with simple, homogeneous dynamics across the entire network. Future work should explore the role of heterogeneous dynamics [36, 155]. Second, the present dynamics do not depend on time-delayed transmission, allowing us to focus only on network structure, but ignoring the role of network geometry in learning [132]. Third, we used computational tractometry to reconstruct connectomes from diffusion weighted imaging, a technique prone to false positives and negatives [33, 83, 140]. Although we implemented steps to focus on highly-reproducible consensus features of these networks, future experiments could be designed around networks reconstructed using invasive methods with greater fidelity, such as tract tracing.

Despite common roots, modern neuroscience and artificial intelligence have followed diverging paths. Technological, analytic and theoretical advances present a unique opportunity for convergence of these two vibrant scientific disciplines. Here, we conceptually bridge neuroscience and artificial intelligence by training brain networks to learn a cognitive task. From the connectomics perspective, this work opens fundamentally new opportunities to discover how cognitive capacity emerges from the links and interactions between brain areas. From the artificial intelligence perspective, reverse-engineering biological brain networks may ultimately generate insights and novel design principles for re-engineering artificial brain-inspired networks and systems.

## METHODS

### Data acquisition

A total of N = 66 healthy young adults (16 females, 25.3 ± 4.9 years old) were scanned at the Department of Radiology, University Hospital Center and University of Lausanne. The scans were performed in 3-Tesla MRI scanner (Trio, Siemens Medical, Germany) using a 32-channel head-coil. The protocol included (1) a magnetization-prepared rapid acquisition gradient echo (MPRAGE) sequence sensitive to white/gray matter contrast (1 mm in-plane resolution, 1.2 mm slice thickness), (2) a diffusion spectrum imaging (DSI) sequence (128 diffusion-weighted volumes and a single b0 volume, maximum b-value 8000 s/mm^2^, 2.2 × 2.2 × 3.0 mm voxel size), and (3) a gradient echo EPI sequence sensitive to BOLD contrast (3.3 mm in-plane resolution and slice thickness with a 0.3 mm gap, TR 1920 ms, resulting in 280 images per participant). Participants were not subject to any overt task demands during the fMRI scan.

### Structural network reconstruction

Grey matter was parcellated into 68 cortical plus 15 subcortical nodes according to the Desikan-Killiany atlas [37]. Cortical regions were then further divided into 1000 approximately equally-sized nodes [26]. Structural connectivity was estimated for individual participants using deterministic streamline tractography. The procedure was implemented in the Connectome Mapping Toolkit [31], initiating 32 streamline propagations per diffusion direction for each white matter voxel. Structural connectivity between pairs of regions was defined as the number of streamlines normalized by the mean length of streamlines and mean surface area of the two regions, termed fiber density [48]. This normalization compensates for the bias toward longer fibers during streamline reconstruction, as well as differences in region size.

To mitigate concerns about inconsistencies in reconstruction of individual participant connectomes [63, 140], as well as the sensitive dependence of network measures on false positives and false negatives [163], we adopted a group-consensus approach [19, 33, 110]. In constructing a consensus adjacency matrix, we sought to preserve (a) the density and (b) the edge length distribution of the individual participants matrices [17, 19, 95]. We first collated the extant edges in the individual participant matrices and binned them according to length. The number of bins was determined heuristically, as the square root of the mean binary density across participants. The most frequently occurring edges were then selected for each bin. If the mean number of edges across participants in a particular bin is equal to *k*, we selected the *k* edges of that length that occur most frequently across participants. To ensure that inter-hemispheric edges are not under-represented, we carried out this procedure separately for inter- and intra-hemispheric edges. The binary density for the final whole-brain matrix was 2.5% on average. The weight associated with each edge was then computed as the mean weight across all participants.

### Reservoir computing

We used reservoir computing to measure the encoding capacity of large-scale brain networks in a memory task. In its simplest form, the reservoir computing architecture consists of an input layer, followed by a recurrent neural network (the *reservoir*), and a readout module, which is typically a linear model. Given an input and a target signal, a learning task consists of approximating the target signal through a linear combination of the states of the readout nodes within the reservoir. The latter are activated by the propagation of the external input signal, introduced to the reservoir through a set of input nodes. Fig. 1a illustrates the paradigm.

Briefly, reservoir computing relies on the two following mechanistic principles: *i)* the complex interactions within the reservoir perform a nonlinear projection of the input into a higher dimensional space; this has the potential to convert nonlinearly separable problems into linearly separable ones; and *ii)* the recurrent nature of the connections within the reservoir endows the network’s states with a temporal memory. These two properties, which also depend on the dynamics of the reservoir, are critical as they constitute some of the basic computational building blocks of more complex tasks. Therefore, if these two properties exist, the reservoir has the potential to serve as a universal function approximator [82].

In the present report, we used a high-resolution human brain connectome reconstructed from diffusion-weighted imaging to constrain the connections within the reservoir. Throughout all experiments, we used all subcortical regions as input nodes, and a subset of cortical regions as readout nodes that were defined based on the Yeo-Krienen intrinsic network partition [141]. Details about network dynamics, training of the network, and the memory task are described below.

#### Reservoir dynamics

The reservoir consists of an artificial recurrent neural network of nonlinear threshold units. Each unit receives one or more inputs and sums them to produce an output. Each input is separately weighted, and the sum is passed through a nonlinear activation function. The dynamics of the reservoir are thus governed by the following discrete-time, firing-rate neuron model:

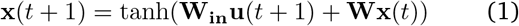

where **x**(*t*) represents the vector of activation states of the nodes inside the reservoir (*reservoir states*). The vector **u**(*t*) = (*u*_1_(*t*)*, , u_K_*(*t*)) is a *K*-dimensional input signal weighted by the input matrix **W_in_**(usually a constant, unless stated otherwise). Finally, the matrix **W** represents the connection weights within the reservoir, i.e., the brain connectivity measures obtained from diffusion MRI data.

#### Stability of reservoir dynamics

We parametrically tuned the dynamics of the reservoir by uniformly scaling the connection weights so that the spectral radius (that is, the modulus of the leading eigen-value) of the connectivity matrix is either below, at, or greater than 1 [118]. Specifically, we first scaled the weights of the connectivity matrix between 0 and 1, and then we divided by its spectral radius. The latter operation transforms the spectral radius of the connectivity matrix to 1. To gradually modified the spectral radius, we multiplied the connectivity matrix by a wide range of *α* values ([0.3, 0.5, 0.7, 0.8, 0.9, 1.0, 1.1, 1.2, 1.3, 1.4, 1.5, 2.0, 2.5, 3.0, 3.5]), being *α* the tuning parameter. The matrix **W** in Eqn. 1 can then be expressed as:

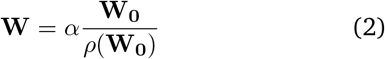

where **W_0_** is the original weighted connectivity matrix scaled between 0 and 1 and *ρ*(**W_0_**) is the spectral radius of **W_0_**. Reservoir dynamics are stable for *α <* 1, and chaotic for *α >* 1. When *α* ≈ 1, dynamics are said to be critical, or at the *edge of chaos*, a transition point between stable and chaotic dynamics [118].

#### Readout module

The role of the readout module is to approximate the target signal **y**(*t*), specific to the task at hand, through a linear combination of the activation states of the output nodes within the reservoir, **x_out_**(*t*). That is:

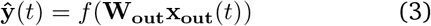

where **W_out_** is the matrix of weights that connect the readout nodes to the readout module. Using the target signal **y**(*t*), these weights are trained in a supervised manner. In principle, learning can be attained using any type of linear regression model. Here, *f* represents the ordinary least squares estimator.

### Intrinsic networks as functional systems

The theoretical paradigm here proposed (Fig. 1a) makes part of a more extensive effort to develop a framework to investigate structure-function relationships across the brain (*function* understood from a computational point of view). Specifically, this framework aims to further our understanding on how functional specialization emerges from the underlying network structure. To investigate the effects of network structure on the emergence of the spectrum of cortical functions, we need a convenient and biologically meaningful way to divide the cortex into its functional systems. Here we used a connectivity-based functional partition known as *intrinsic networks*.

Intrinsic networks, often known as *resting-state networks* in the cognitive neuroscience literature, consist of brain regions that exhibit high levels of synchronized BOLD activity [159]. It has been hypothesized that the organization of the brain into these segregated systems or modules supports functional specialization [159]. These internally coherent and co-fluctuating networks are associated with individual differences in task activation and behaviour, including perception, cognition and action [29, 89]. In the present study, we applied the intrinsic network partition derived by Yeo, Krienen and colleagues [141]. Briefly, the partition was obtained by applying a clustering algorithm to voxel-level functional connectivity estimated from resting-state functional MRI data. These networks have been consistently replicated in multiple studies using different data acquisitions, anatomical parcellations and analysis techniques [32].

### Memory task

Using the experimental paradigm described above, we estimated the encoding capacity of each intrinsic network in a memory task. In this task, the readout module is trained to reproduce a time-delayed random input signal at various lags. In other words, **y**(*t*) = **u**(*t − τ*), where **y**(*t*) is the target signal, **u**(*t*) is the input signal, and *τ* is the parameter that controls for the amount of memory required by the task.

We generated 4100 points of a uniformly distributed input signal (**u**(*t*) ∼ Uniform(−1, 1)). We introduced this signal to the reservoir through a set of input nodes corresponding to all 15 subcortical regions. We then recorded, for the same length of time, the activation states of the readout nodes (these activation states are represented by **X_out_** in Eqn. 3). We used 2050 time points of the recorded activation states to train a linear regression model to reproduce the same input signal at different time lags (see *Readout module* for details); *τ* was monotonically increased in one-size steps in the range [1, 16]. The remaining 2050 time points were used to test the performance. For every *τ*, a performance score was computed as the absolute value of the Pearson correlation between the target **y**(*t*) and the predicted signal 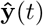. Memory capacity (MC) was then estimated as the sum of the performance score across all time delays:

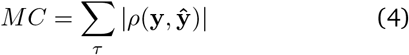

The magnitude of MC is proportional to the ability of the reservoir’s activation states to encode both past and present input stimuli.

The procedure above was repeated for every value of the parameter *α* (see *Stability of reservoir dynamics* for details), and for every intrinsic network. In this way, the net result is a memory capacity per intrinsic network as a function of the *α*.

### Bootstrap resampling

We bootstrapped individual connectivity matrices to provide a reliable estimate for the memory capacity of each intrinsic network. Specifically, we generated 1000 (sub)samples by randomly selecting, without replacement, 40 out of 66 individual connectivity matrices. For each drawn subset, we generated a group-consensus structural connectivity matrix (see *Structural network reconstruction* for details), and we applied the procedure described in the previous section. In this way, we constructed a distribution for the memory capacity of each functional system at every value of the parameter *α* (Fig. 1c).

### Null models

We benchmarked the memory capacity of intrinsic networks against two different types of null models. The first null model evaluates the extent to which memory capacity measures depend on the underlying network connectivity. To do so, we used the method proposed in [86], available in the Python version of the Brain Connectivity Toolbox (https://github.com/aestrivex/bctpy; [112]). This method systematically destroys network topology by randomly swapping pairs of edges (10 swaps per edge), while preserving network size, degree distribution, density, and node-level, intrinsic-network assignment (Fig. 2b). This method guarantees the connectedness of the resulting rewired network, that is, every node can be reached by every other node in the network.

The second null model assesses whether the partition of the connectome into seven intrinsic networks is relevant for the performance in the memory task. We used a spatially-constrained, label-permutation null model that randomly permutes intrinsic network labels while preserving their spatial embedding and autocorrelation (Fig. 2c; [1, 85]). This method creates a surface-based representation of our parcellations by applying the Lausanne atlas to the FreeSurfer *fsaverage* surface using files obtained from the Connectome Mapper toolkit (https://github.com/LTS5/cmp) [31]. The *fsaverage* surface is then projected onto a sphere to define spatial coordinates for each parcel; vertices on the sphere are assigned based on the closest distance to the center-of-mass of each parcel. New spatial coordinates are generated by applying randomly-sampled rotations. Finally, node-labels are then reassigned based on that of the closest resulting parcel. Importantly, this procedure was performed at the parcel resolution rather than the vertex resolution to ensure that parcel sizes were not changed during the rotation and reassignment procedure. This method is available in the module *stats* of the Net-neurotools python package (https://netneurotools.readthedocs.io/en/latest/index.html).

These models were applied on a consensus structural connectivity matrix reconstructed from the individual matrices of all 66 subjects. A null distribution with 1000 iterations was built for each model. We used the Wilcoxon-Mann-Whitney two-sample rank-sum test to assess the statistical significance of the human connectome’s memory capacity estimates against the null models.

### Wiring cost

Euclidean distance was used as a proxy for white matter tract length. We estimated the distance between every pair of brain regions that are physically connected by a structural connection. The total wiring cost for the network was then estimated as the sum of the connection lengths weighted by the connectivity values; the latter are proportional to the number of streamlines between brain regions. Same conclusions can be drawn when the mean or median of the connection length distribution are used instead (not shown in the present report).

### Encoding and decoding capacity

Broadly speaking, any communication scheme is characterized by the presence of two agents: the *encoder*, whose role is to *encode* and *send* a message, and the *decoder*, who is in charge of *receiving* and *decoding* the message. In such scheme, the capacity of the encoder agent to cipher – or encode – the message is bound to the information it *sends*, and the capacity of the decoder agent to decipher – or decode – the message is bound to the information it *receives*. Following a similar logic, we operationalized the *encoding* and *decoding* capacity of an intrinsic network as the memory capacity of the information *sent* and *received* by the network, respectively. This translates into two different experimental set ups, namely *encoding* and *decoding*, which conceptually diverge in the role played by the network (i.e., encoder or decoder), and pragmatically differ in the way read-out nodes are selected. For *encoding*, the signals generated by the nodes of the network are used as a proxy for the information *sent* by the network. For *decoding*, the signals generated by the directly connected neighbour nodes of the network are used as a proxy for the information *received* by the network.

#### Encoding - decoding: interpretation

To quantify how temporal information content is transformed as it flows through a particular functional network, we considered the difference between its encoding and decoding capacity (*encoding decoding*). A large difference between encoding and decoding indicates that the memory content of the information traveling from the rest of the brain to an intrinsic network has undergone a significant transformation. If the difference between encoding and decoding is positive (*encoding > decoding*; good encoder), this suggests that the interaction between intrinsic network connectivity and dynamics enhances the memory content of the signal within the network to perform the task at hand. Conversely, a negative difference between encoding and decoding (*encoding < decoding*; good decoder) suggests that the interaction between extrinsic network connectivity and dynamics rather favours the memory content of the signals coming to the network. Summing up, the magnitude of the difference between the encoding and decoding capacity of a network is proportional to the memory content gained or lost by the signal as information propagates through the network. The sign of the difference tells us the direction in which this transformation occurred, in other words, whether the signal gained (positive) or lost (negative) temporal content as it travels from the rest of the brain to a particular functional network.

#### Methodological details

The selection of the readout nodes in the experimental set up is what ultimately determines the *coding* paradigm. In the encoding set up, readout nodes correspond to the nodes within the network, whereas in the decoding setup, readout nodes correspond to the direct neighbours of the network. In the latter case, because the number of neighbour regions (i.e., directly connected nodes that do not belong to the network) is in all cases higher than the amount of regions within each intrinsic network, these differences should be accounted for when estimating their encoding and decoding capabilities. To do so, we first derived the between-network connectivity profile of every node in the network under examination. Based on these node-level connectivity profiles, we estimated the proportion of connections from every other network (within-network connections were excluded). Next, we randomly draw as many neighbour regions as nodes within the network in such a way that the proportions previously estimated were preserved. The selected nodes were then used as output nodes in the decoding experimental set up. To avoid biases towards a particular set of output nodes, the same process was repeated 100 times to get a better estimate of the decoding capacity of every intrinsic network.

It is important to note, however, that the metric here proposed, i.e., the difference between encoding and decoding capacity, assumes that signals propagate in a particular direction. Because MRI-based structural networks do not contain information about the directionality of anatomical projections [70, 90], our network model is undirected and thus considers that information travels bidirectionally. Therefore, interpretation of the results should be done with respect to this limitation.

### Graph network properties

We explored the effects of the interaction between network topology and dynamics on the memory capacity of intrinsic networks. To do so, we correlated local and global network topological attributes with the memory capacity of the brain’s intrinsic networks. Local topological attributes included: (1) node strength, (2) clustering coefficient, (3) node-betweenness centrality and (4) participation coefficient. These local features were first estimated at the node level using the weighted adjacency matrix, and then averaged within each intrinsic network. Global topological features included: (1) characteristic path length, (2) transitivity and (3) modularity. Global features were estimated at the network level, and then correlated with the average memory capacity across intrinsic networks. To calculate all graph topological metrics we used a Python version of the Brain Connectivity Toolbox (https://github.com/aestrivex/bctpy; [112]) Definitions of these topological metrics can be found below.

- *Node strength:* in undirected networks, node strength is defined as the sum of the weights of the connections that go from a node to every other node in the network.
- *Clustering coefficient:* the local clustering of a node is the proportion of connections between the nodes within its neighbourhood divided by the total number of possible connections between them [100].
- *Node betweenness centrality:* the betweenness centrality of a node is the fraction of all shortest paths in the network that contain the node. Nodes with a high value of betweeness centrality participate in a large number of shortest paths [21].
- *Participation coefficient:* the participation coefficient is a measure of the distribution of a node’s connections among the communities of the network. If the participation coefficient is 0, the connections of a node are entirely restricted to its community. The closer the participation coefficient is to 1, the more evenly distributed are the connections of the node among all communities [46]. Mathematically, the participation coefficient *P* of node *i* is given by:

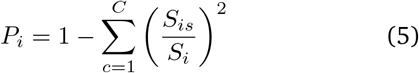

where *S_is_* corresponds to the sum of the weights of the connections from node *i* to nodes in community *c*, *S_i_* is the strength of node *i*, and *C* is the total number of communities. This measure requires a previously established community structure as input. Here, we estimated the participation coefficient of individual brain regions using the proposed intrinsic-network partition.
- *Characteristic path length:* the characteristic path length is a measure of efficiency and is defined as the average shortest path length of the network. The distance matrix on which shortest paths are computed must be a connection-length matrix, typically obtained via a mapping from weight to length. Here we used the inverse of the connection-weight matrix as the distance matrix. In this way, strong (weak) connections are naturally interpreted as shorter (longer) distances [38, 158].
- *Transitivity:* a network’s transitivity is the ratio of triangles to triplets (open and closed) in the network. A triplet consists of three nodes that are connected by either two (open) or three (closed) undirected connections [100, 112, 158].
- *Modularity:* is a measure that relates the number of within-network connections to all connections in the network; it quantifies the strength of segregation into distinct networks. The higher the modularity, the stronger is the segregation/separation between modules [44, 76, 107]. Mathematically, the modularity *Q* of a network can be expressed as:

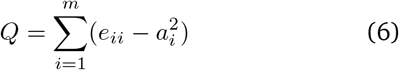

where *e_ii_* is the fraction of all connections that link two nodes within module *i*, *a_i_* is the fraction of connections that connect a node in module *i* to any other node, and *m* is the total number of modules. As with the participation coefficient, this measure requires a previously established community structure as input. We used the intrinsic-network partition for this purpose.

## Code availability

The Python repository used for the simulations of the reservoir and the implementation of the memory capacity task, as well as for the generation of the null network models is available on GitHub (https://github.com/estefanysuarez) and is built on top of the following open-source Python packages: Numpy [51, 99, 154], Scipy [153], Pandas [88], Scikitlearn [102], Bctpy (https://github.com/aestrivex/bctpy) [112], NetworkX [47], Netneurotools https://netneurotools.readthedocs.io/en/latest/, Mat-plotlib [59], and Seaborn [157].

## ACKNOWLEDGEMENTS

We thank Ross Markello, Bertha Vazquez-Rodriguez, Golia Shafiei, Vincent Bazinet, Justine Hansen and ZhenQi Liu for insightful comments on the manuscript. BM acknowledges support from the Natural Sciences and Engineering Research Council of Canada (NSERC Discovery Grant RGPIN #017-04265), from the Canada Research Chairs Program, and from the Canada First Research Excellence Fund, awarded to McGill University for the Healthy Brains for Healthy Lives initiative. GL [NSERC Discovery Grant (RGPIN-2018-04821) and FRQS Research Scholar Award, Junior 1 (LAJGU0401-253188)]. BAR acknowledges support from NSERC (NSERC Discovery Grant RGPIN 2020-05105) and CIFAR (Learning in Machines and Brains Program). ES acknowledges support from the FRQNT Strategic Clusters Program (2020-RS4-265502 - Centre UNIQUE - Union Neurosciences and Artificial Intelligence - Quebec).

---

**Figure S1.**
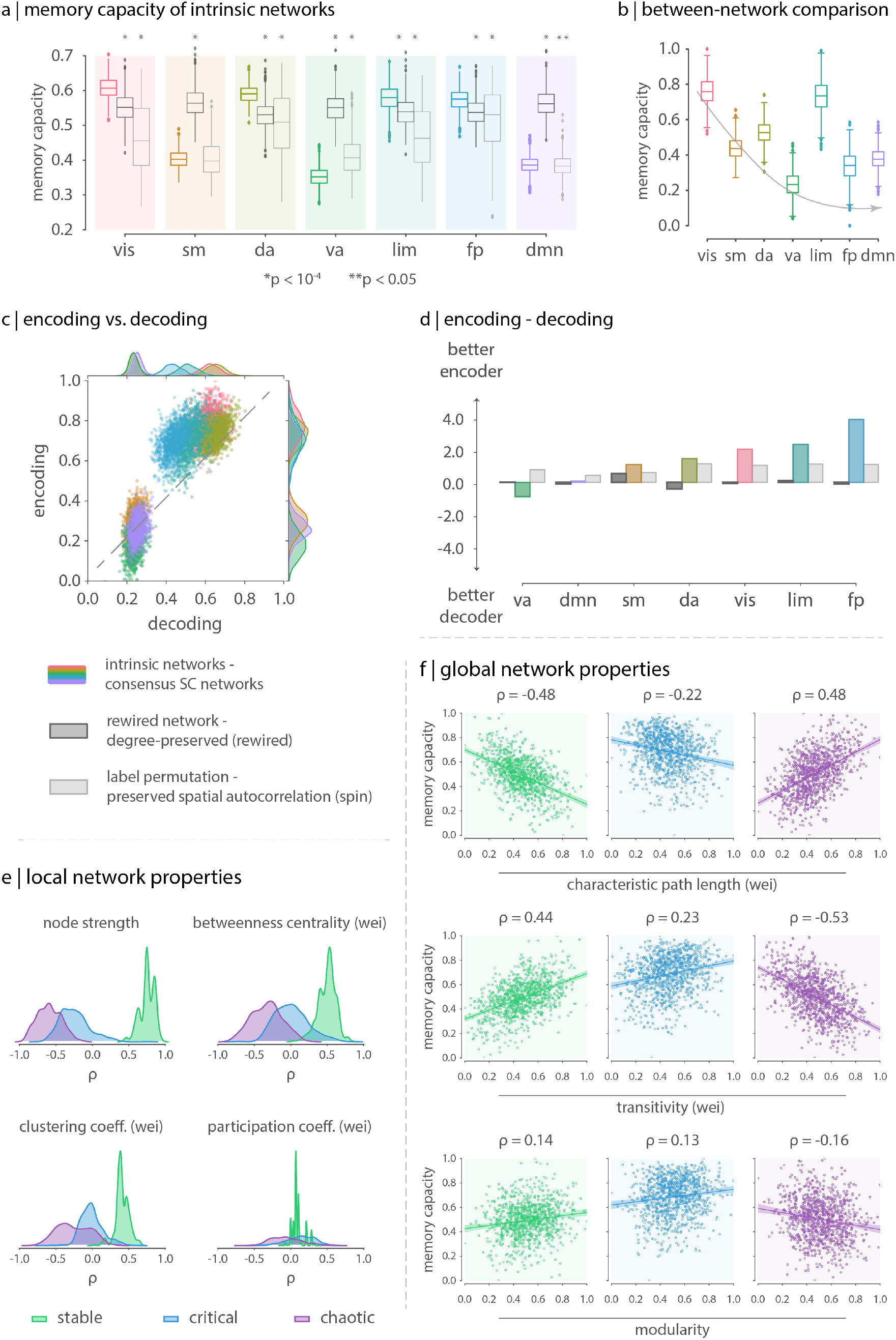
A chaotic state closer to critical dynamics. Results in the chaotic regime were replicated for alpha values in the range *α* = [1.1, 1.2, 1.3, 1.4, 1.5]. (a) Memory capacity of intrinsic networks compared to the rewired and label-permutation models. (b) Between network comparison of memory capacity estimates after the removal (through linear regression) of relative connection density effects. (c) Encoding vs decoding capacity of intrinsic networks. Encoding and decoding capacity values were scaled between 0 and 1 according to the maximum and minimum values in the chaotic regime. (d) Information transfer of intrinsic networks. Relationship between memory capacity and (e) local and (f) global network properties.

**Figure S2.**
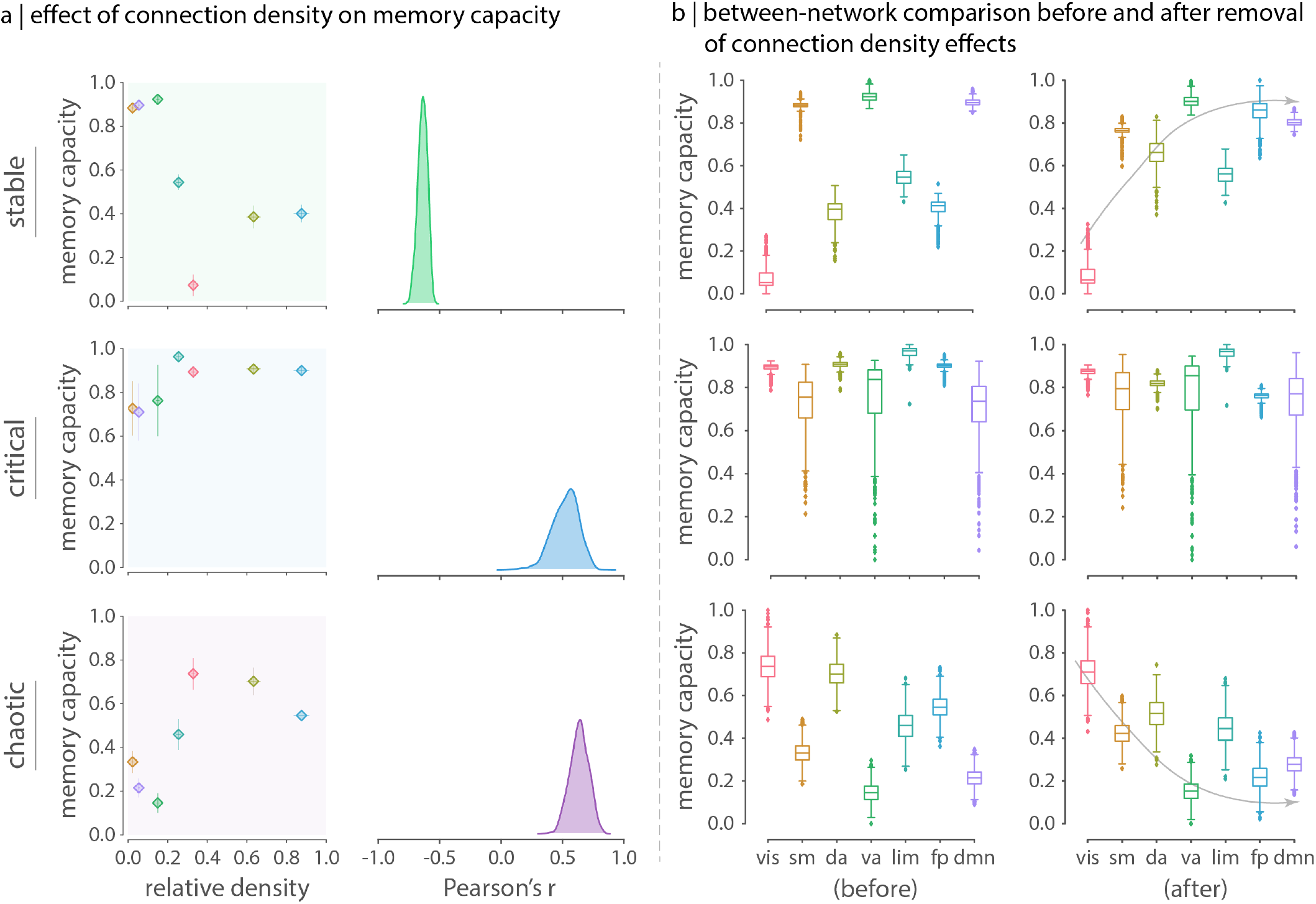
Effects of relative density on the memory capacity of intrinsic networks. The relative density of an intrinsic network is defined as the ration of the number of internal connections to the total number of possible connections within the intrinsic network. (a) The left column shows the relationship between memory capacity and relative density across dynamical regimes (markers represent the mean values across bootstrapped consensus matrices and error bars correspond to their standard deviation). The right column shows the distribution (across bootstrapped consensus matrices) of the Pearson’s correlation between the memory capacity of intrinsic networks and their relative density. (b) Memory capacity of intrinsic networks before (left column) and after (right column) removing the effects of relative density. Given the high average values of the Pearson’s correlation between memory capacity and relative density, relative density effects were regressed out using a linear model.

**Figure S3.**
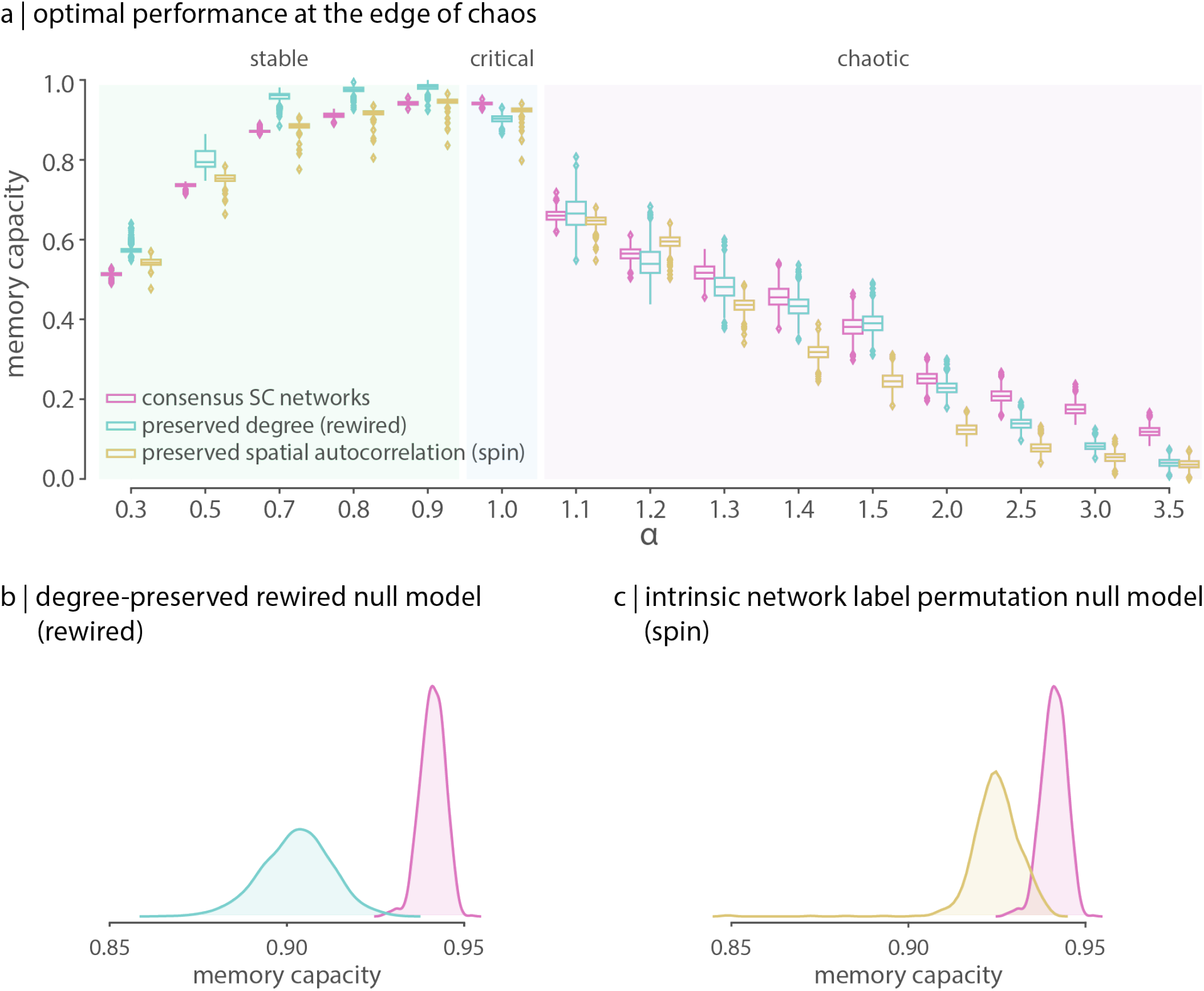
Memory capacity of the brain - replication using a more biologically-plausible set of input nodes. Analyses were replicated considering thalamic subcortical regions as input nodes. (a) Distribution (across bootstrapped consensus matrices) of the memory capacity of the human connectome (magenta) as a function of the dynamics. Similar to the experimental set up where all subcortical regions are used as input nodes (Fig.2), the estimated memory capacity of the human connectome is maximum and optimal at the edge of chaos (*α* = 1). (b) Distribution of the memory capacity of the empirical network (magenta; median = 0.94) vs. the rewired network model (cyan; median = 0.90, *P_rewired_* = 0.0 and effect size = 99% two-tailed, Wilcoxon-Mann-Whitney two-sample rank-sum test) at the critical state. (c) Distribution of the memory capacity of the empirical network (magenta) vs. the label-permutation model (yellow; median = 0.92, *P_spin_* = 0.0 and effect size = 99% two-tailed, Wilcoxon-Mann-Whitney two-sample rank-sum test) at the critical state.

**Figure S4.**
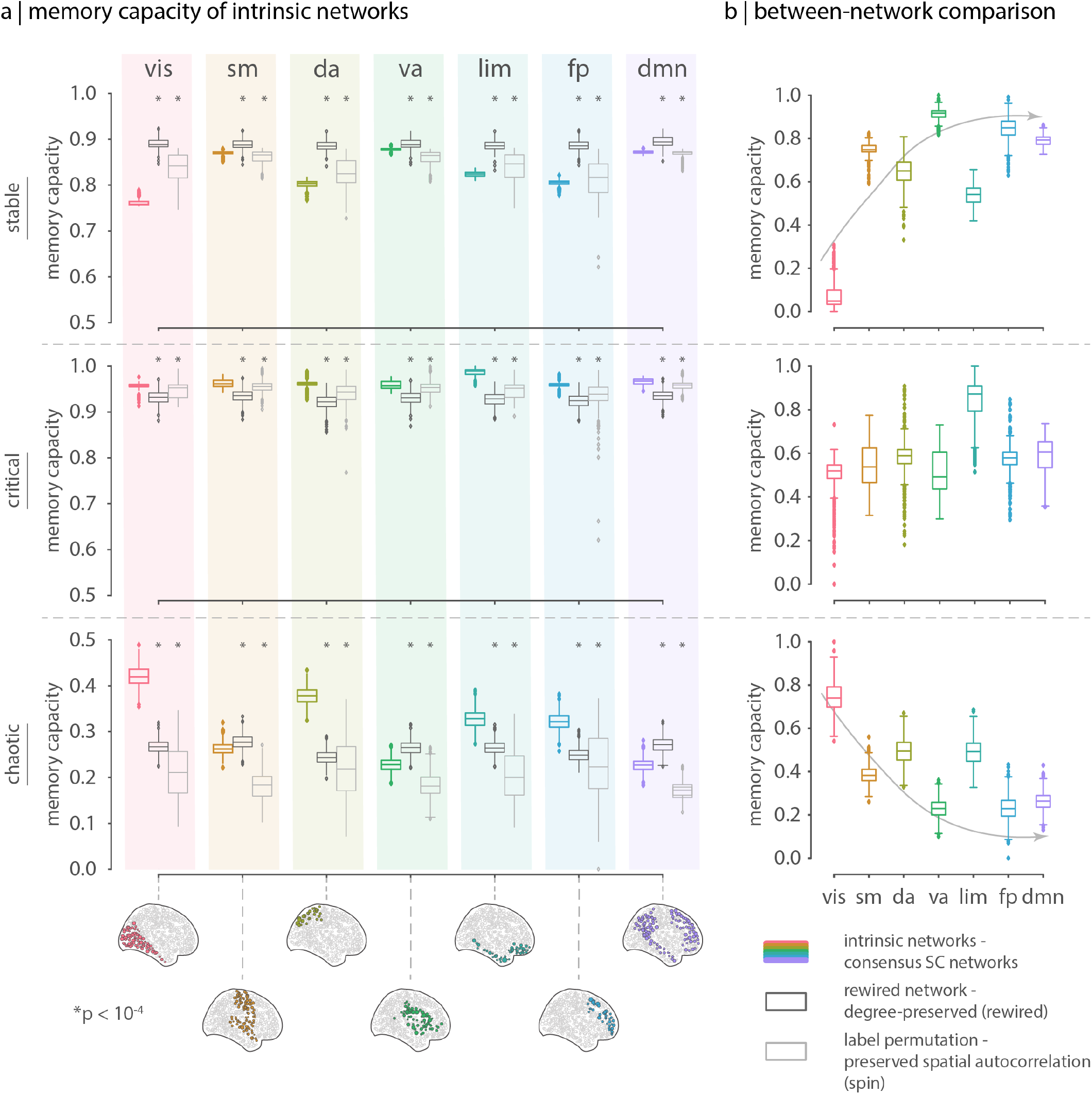
Memory capacity of intrinsic networks - replication using a more biologically-plausible set of input nodes. Analyses were replicated considering thalamic subcortical regions as input nodes. (a) Memory capacity of intrinsic networks across dynamical regimes. At the critical state, memory capacity estimates on the empirical network are consistently and significantly higher and optimal across functional systems, compared to the rewired and spatially-constrained null models. (b) Comparison across intrinsic networks, after linearly regressing out the effect of relative connection density on memory capacity estimates. When compared to each other, memory capacity is in general quite homogeneous across functional systems in the critical regime, except for the limbic system, for which memory capacity is the highest, maybe reflecting an anatomically-mediated predisposition for memory encoding. In the stable and chaotic regimes, on the other hand, the axis along which memory capacity estimates fluctuate broadly resembles the unimodal-transmodal hierarchy, differentiating again somatosensory from higher-order association areas.

**Figure S5.**
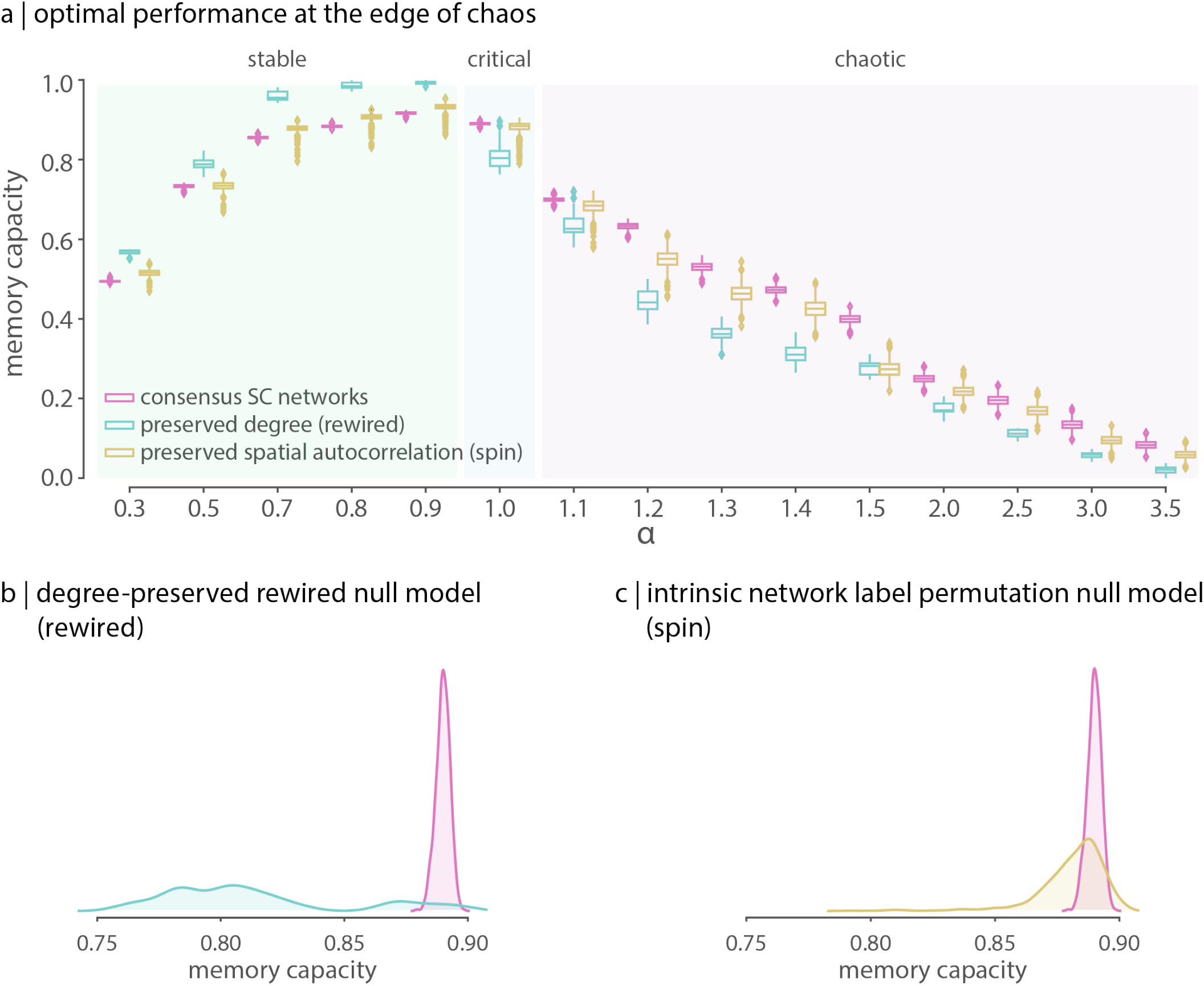
Memory capacity of the brain - replication at a lower scale. Analyses were replicated using a lower resolution parcellation with 448 cortical and 15 subcortical regions. (a) Distribution (across bootstrapped consensus matrices) of the memory capacity of the human connectome (magenta) as a function of the dynamics. Similar to the high-resolution network, the estimated memory capacity of the human connectome is maximum and optimal at the edge of chaos (*α* = 1). (b) Distribution of the memory capacity of the empirical network (magenta; median = 0.89) vs. the rewired network model (cyan; median = 0.80, *P_rewired_* = 10^−4^ and effect size = 95% two-tailed, Wilcoxon-Mann-Whitney two-sample rank-sum test) at the critical state. (c) Distribution of the memory capacity of the empirical network (magenta) vs. the label-permutation model (yellow; median = 0.88, *P_spin_* = 10^−4^ and effect size = 75% two-tailed, Wilcoxon-Mann-Whitney two-sample rank-sum test) at the critical state.

**Figure S6.**
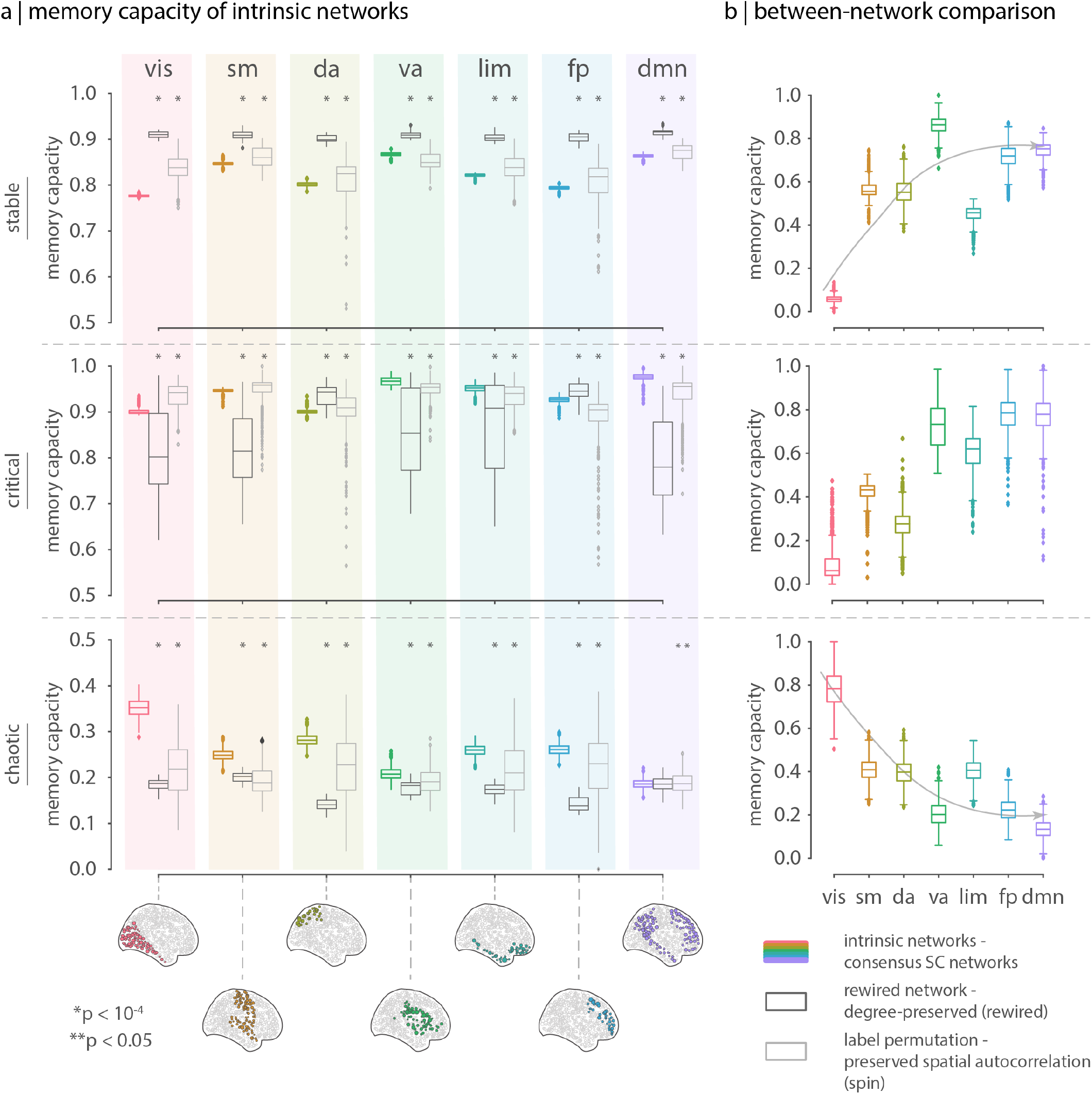
Memory capacity of intrinsic networks - replication at a lower scale. Analyses were replicated using a lower resolution parcellation (448 cortical and 15 subcortical regions). (a) Memory capacity of intrinsic networks across dynamical regimes. In contrast to the high resolution network, at the critical state, memory capacity estimates on the empirical network are significantly different but not consistently higher across all functional systems, compared to the rewired and the spatially-constrained label permutation models. (b) Comparison across intrinsic networks, after linearly regressing out the effect of relative connection density on memory capacity estimates. At the critical state, contrary to what occurs in the high resolution network (in which there are small differences in memory capacity across functional systems), there is a higher differentiation across intrinsic networks’ memory capacity, and a more prominent difference between lower somatosensory and higher association areas. The same type of differentiation is present in the stable and chaotic regimes, but in opposing directions with respect to each other. In the stable regime, memory capacity estimates follow the same trend observed in the critical state, in which somatosensory areas are on average lower compared to higher association areas. The opposite is observed in the chaotic regime: lower somatosensory areas present on average higher memory capacity compared to higher association areas.

